# Assessing the Dynamics and Control of Droplet- and Aerosol-Transmitted Influenza Using an Indoor Positioning System

**DOI:** 10.1101/130658

**Authors:** Timo Smieszek, Gianrocco Lazzari, Marcel Salathé

**Affiliations:** Modelling and Economics Unit, National Infection Service, Public Health England, London, United Kingdom; MRC Centre for Outbreak Analysis and Modelling, Department of Infectious Disease Epidemiology, Imperial College School of Public Health, London, United Kingdom; Center for Infectious Disease Dynamics, The Pennsylvania State University, University Park, USA; Global Health Institute, School of Life Sciences, Ecole Polytechnique Fédérale de Lausanne (EPFL), Lausanne, Switzerland

## Abstract

There is increasing evidence that aerosol transmission is a major contributor to the spread of influenza. Despite this, virtually all studies assessing the dynamics and control of influenza assume that it is transmitted solely through direct contact and large droplets, requiring close physical proximity. Here, we use wireless sensors to measure simultaneously both the location and close proximity contacts in the population of a US high school. This dataset, highly resolved in space and time, allows us to model both droplet and aerosol transmission either in isolation or in combination. In particular, it allows us to computationally assess the effectiveness of overlooked mitigation strategies such as improved ventilation that are available in the case of aerosol transmission. While the effects of the type of transmission on disease outbreak dynamics appear to be weak, we find that good ventilation could be as effective in mitigating outbreaks as vaccinating the majority of the population. In simulations using empirical transmission levels observed in households, we find that bringing ventilation to recommended levels has the same mitigating effect as a vaccination coverage of 50% to 60%. Our results therefore suggest that improvements of ventilation in public spaces could be an important and easy-to-implement strategy supplementing vaccination efforts for effective control of influenza spread.

## Introduction

Despite extensive clinical experience and decades of research on influenza, it is still not fully understood how influenza is transmitted among humans. Traditionally, influenza transmission has been assumed to occur through the air, physical contact between humans, and by touching contaminated surfaces (i.e., fomites). Airborne transmission can occur in two ways: either through relatively large particles of respiratory fluid (droplets) or through smaller such particles that can remain aerosolized (droplet nuclei). As larger droplets are pulled to the ground by gravity quickly, droplet transmission requires close physical proximity between infected and susceptible individuals, whereas aerosolized transmission can occur over larger distances and does not necessarily require that infected and susceptible individuals are at the same location at the same time.

Until recently, close contact transmission was considered to be the dominant transmission pathway, largely because the evidence to support the importance of transmission through aerosols was mixed^1, 2^. However, the question of the importance of the various transmission routes has received renewed attention recently, and multiple studies have in the past few years provided evidence for the importance of aerosol transmission^3–9^. There is increasing evidence from experiments with mammalian hosts that airborne transmission is much more efficient than fomite transmission (see e.g.^5^). Data from randomized controlled trials of hand hygiene and surgical face masks in households provided evidence that aerosol transmission accounts for half of all transmission events^8, 9^. A nosocomial influenza outbreak with subsequent airflow analysis provided further evidence for the important role of aerosol transmission^6^. An experimental laboratory study using a patient examination room containing a coughing manikin provided further support for aerosol transmission^7^. A recent study with outpatients who tested positive for influenza A virus demonstrated that 53% and 42% produced aerosol particles containing viable influenza A virus during coughing and exhalation, respectively^10^. Last but not least, a mathematical model of influenza transmission within a household has suggested that the aerosol transmission route may not only be important, but indeed dominant^3^.

Given the increasing evidence supporting an important role of aerosol transmission, it is prudent to revisit our expectations on disease dynamics of influenza outbreaks, and the best measures to control the spread of influenza. To address this issue, we use a high-resolution dataset of a medium-sized US high school, where both individual’ s indoor positions and close proximity contacts to others were measured using wireless sensor network technology (described in detail in^11^ and in the Methods). Computational simulations on this dataset, parametrized with empirical transmission levels observed in households, suggest that bringing ventilation levels to recommended values has the same effect on final outbreaks sizes as vaccinating between 50 and 60% of the population. Given the ongoing public health challenge of increasing or maintaining high influenza vaccination rates^12^, these results suggest that proper room ventilation is a highly underestimated control measure for effective influenza control.

## Results

We first investigate the dynamics of influenza spread in three different transmission models, namely a droplet-based, an aerosol-based, and a combined droplet-aerosol-based model (for a schematic explanation of these transmission models, please see fig. 1). These three models were chosen to compare the two extreme situations (droplet-only, and aerosol-only) as well as an intermediate situation where the two transmission modes are equally relevant (see for example^8, 9^). Simulations of influenza outbreaks were based on an SEIR model run on a high-resolution contact network collected at a US high school (^11^) using wireless sensor network technology. In addition, we used the location information of each individual obtained using the same technology. In order to simulate partial or full aerosol transmission, we combined this data with building data from the school in order to compute the relevant infection probabilities due to aerosol transmission as per eq. 2. The entire model, and the data sources, are described in full detail in the Methods. Figure 2 shows an example of quanta concentrations in multiple classrooms during a day.

**Figure 1.**
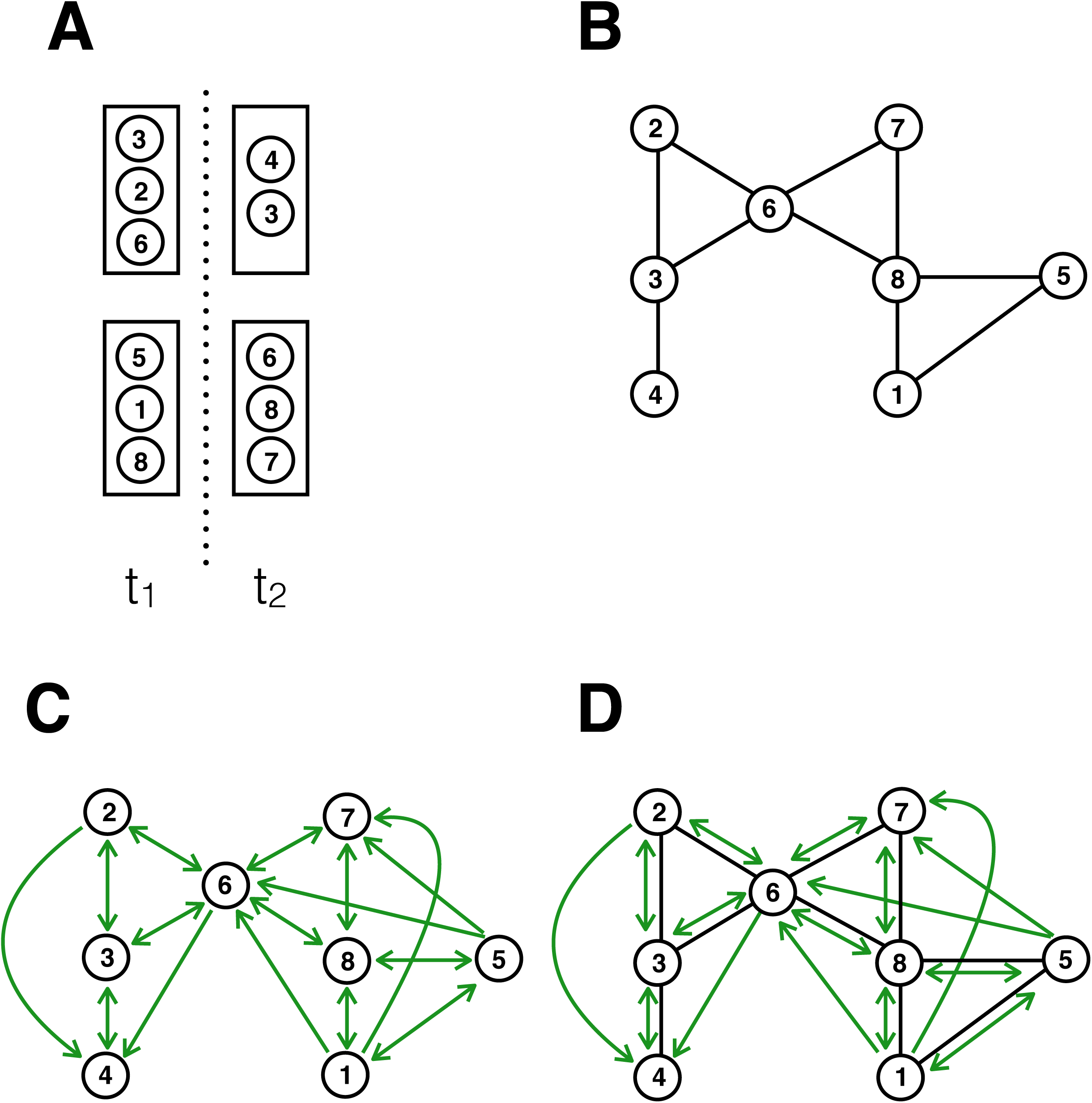
Simplified scheme of transmission routes. Different sets of individuals in the school can occupy the same rooms, at different times t1 and t2 (A). In these rooms, individuals may be in close proximity to one another. For visual simplicity, we assume here that all individuals in the same room at the same time are in close proximity. In the aerosol model, infected individuals can shed infectious material while in the room, which in turn may infect those individuals in the room concurrently or later on. The network of possible transmission pathways will therefore look different depending on the transmission routes. Based on the spatio-temporal pattern shown in panel (A), panel (B) shows the network of pure droplet transmission; panel (C) shows the network of pure aerosol transmission; and panel (D) shows the network of droplet and aerosol transmission combined. The edges for droplet transmission are always bidirectional, hence no arrows are shown. The edges for aerosol transmission may be unidirectional due to the temporal delay of virus shedding and virus uptake, hence arrows are shown.

**Figure 2.**
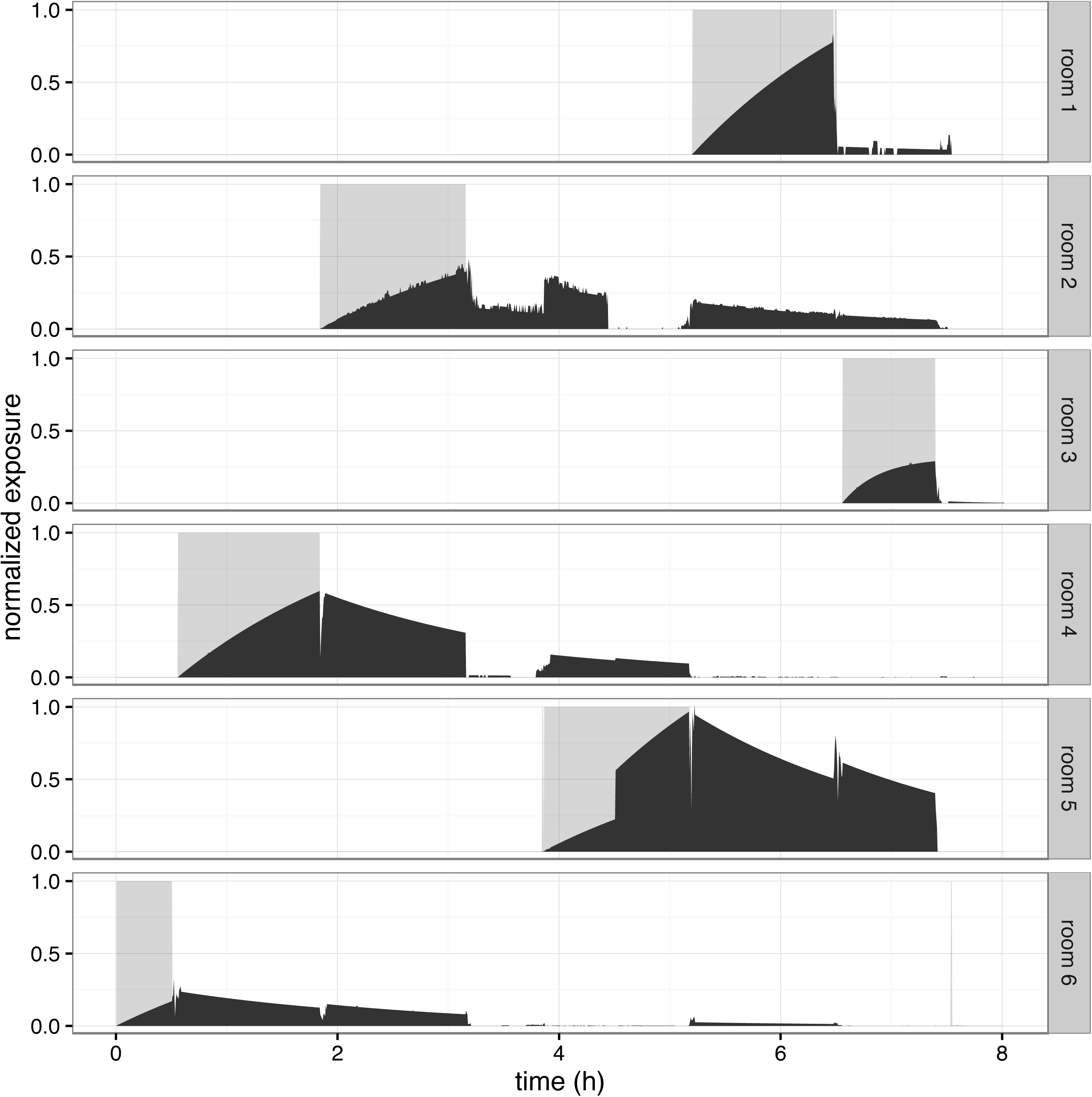
Presence of one individual in different rooms (gray area) and other people’ s exposure to infectious material shed by the individual in the respective rooms (black bars) across 8 hours; the scale of the x-axis is hours, the scale of the y-axis is exposure in [0 - 0.02] quanta. Exposure levels depend on the number of exposed individuals in the room following deposition of infectious material, as well as air change rates.

For all three transmission models, we measure three epidemiological quantities: the final size of an outbreak (number of recovered *r*(*t*) individuals after a simulation run), the total duration of an outbreak (time until there are no more exposed *e*(*t*) or infected *i*(*t*) individuals, respectively), and the time to reach the peak of the outbreak, i.e. to reach the maximal prevalence. In addition, in order to be able to put these quantities in context, we also measure *R*_0_ for the three transmission models. In an individual-based model like the one used here, measuring *R*_0_ is straightforward using the droplet-based transmission model, because one can directly track which individual infected which. However, in the case of partial or full aerosol-mediated transmission where infection is mediated by the air in a room, this tracking is harder because one would need to computationally keep track of the individual sources of infectious aerosol particles. We thus measure an alternative but similar quantity, namely the number of all cases infected during the initial time period until the index case recovers. We call this quantity 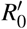. Theoretically, there is a chance that this overestimates the true *R*_0_ by including third generation cases that were not infected by the index case, but given the values for the incubation period and the recovery rate, this is rather rare.

In fig. 3, we can observe that increasing the relative importance of aerosol-based transmission has no major effect overall on disease dynamics. We note that outbreak sizes in the pure droplet model are slightly increased, and the time to outbreak peak is slightly increased. However, the small increase in outbreak size is also reflected in a small increase of 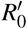 (see fig. 3-D). In summary, our results indicate that one should not necessarily expect influenza disease dynamics to be different when taking aerosol-based transmission into account.

**Figure 3.**
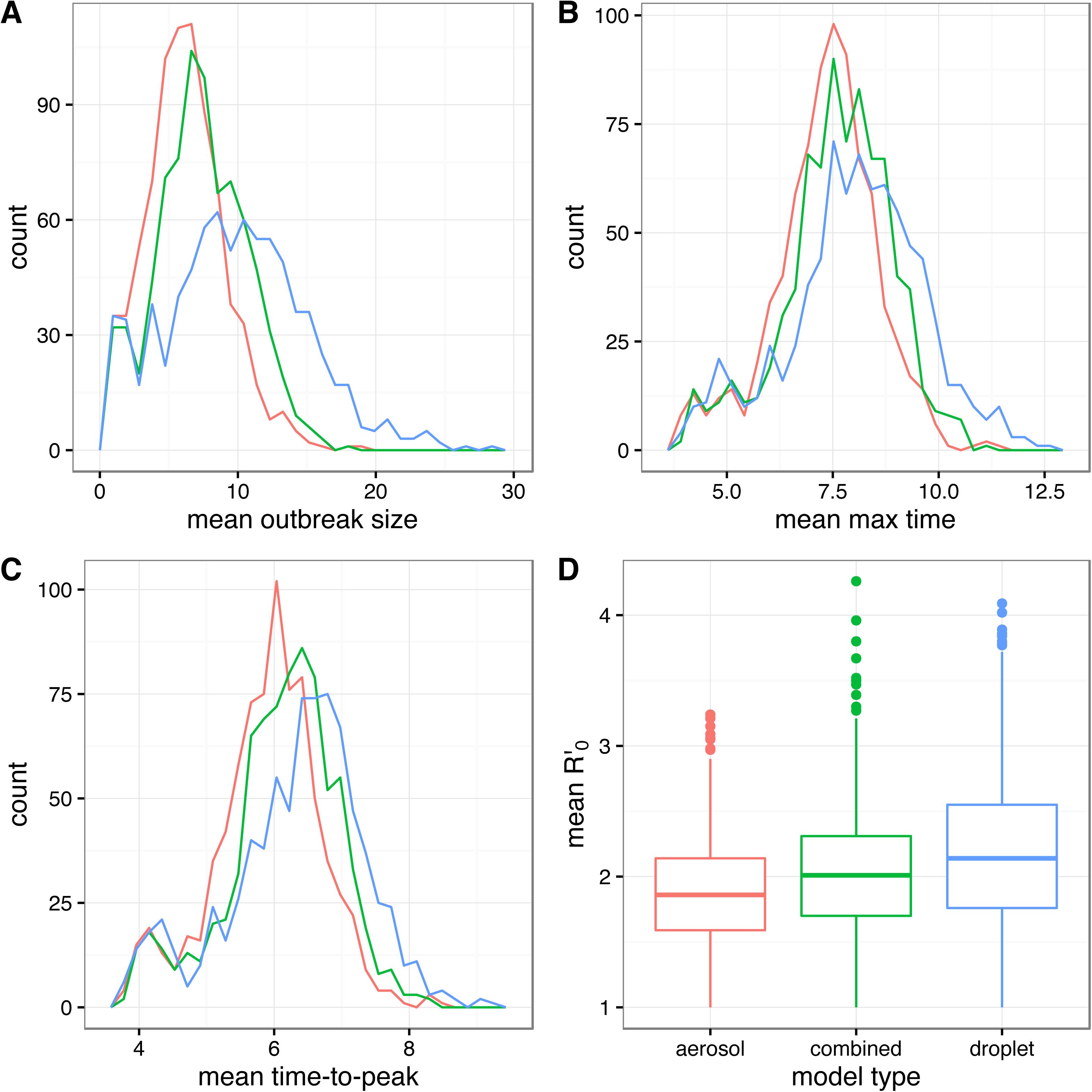
Frequency distributions of mean final size (A), mean duration of outbreak (B) mean time to reach the maximum prevalence (C), and mean 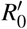, for the three different transmission models. For each transmission model, the values are based on 78,900 simulations where each individual served as index case in 100 independent simulation runs.

In pure droplet-based transmission models, vaccination is a powerful strategy to mitigate the spread of an infectious disease. When transmission can also be aerosol-based, increasing ventilation is an additional way to curb the spread of disease. We therefore compared the effect of ventilation to traditional vaccination strategies. According to the American Society of Heating, Refrigerating and Air Conditioning Engineers (ASHRAE)^13^, a good ventilation in classrooms corresponds to 3 air changes per hour. Most classrooms, however, have poor ventilation at rates around 0.5 air changes per hour (see Methods for more details).

In the pure aerosol-based model, bringing all rooms to the recommended ventilation rate would almost completely eliminate the chance of an outbreak. This corresponds to the same effect of complete vaccination coverage in the case of poor ventilation rates, as shown in fig. 4-B. In the combined droplet-aerosol scenario, improvements of ventilation still results in a significant decrease of outbreak sizes. In particular, fig. 4-A shows that in the combined droplet-aerosol model, a good ventilation would have a similar effect to a 50-60 % vaccination coverage in the poor ventilation scenario.

**Figure 4.**
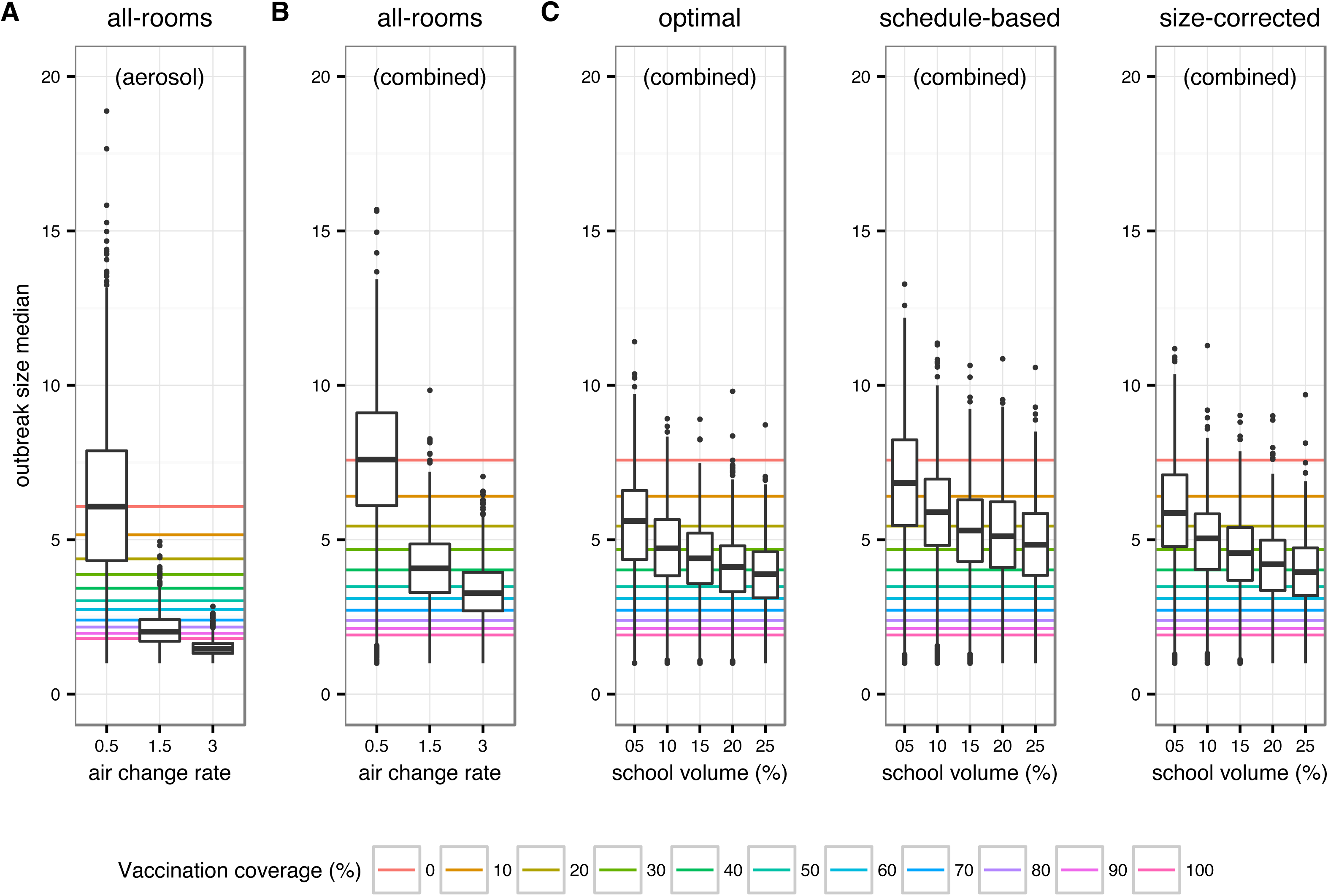
Comparison of the effect of ventilation (boxplots) and vaccination coverage (horizontal lines) on outbreak size. Colors refer to different vaccination coverages. Results are reported for aerosol-based (A) and combined transmission models (B, C). For (A) and (B), air change rate varies from 0.5 to 3.0 changes/h, while (C) assumes an air change rate of 0.5 changes/h. (C) Comparison of partially improved ventilation strategies (boxplots) in the school. Classrooms for improved ventilation were selected by ranking them according to the amount of inhaled infectious particles (optimal), their occupancy according to the school roster (schedule-based), or occupancy corrected by room size (size-corrected). Effect of vaccination coverage is also reported (median outbreak sizes under corresponding vaccination coverages, reported as horizontal lines).

In practice, upgrading the ventilation system of an entire school campus to the rates proposed by ASHRAE will often be challenging due to limited resources. We therefore asked how strong the mitigating effect would be of upgrading the ventilation of only a fraction of all rooms. We also asked how one would identify the optimal set of rooms for mitigation purposes. The room selection strategies we explore are (i) *optimal*, using all available information from simulation runs to identify rooms with the highest cumulative exposure, i.e., where most susceptibles will be exposed or where doses are highest in a typical simulation run; (ii) *schedule-based*, using a school’ s official roster to identify rooms with the highest cumulative occupancy throughout a school day; and (iii) *size-corrected*, which is similar to the schedule-based approach, but the total occupancy was divided by the volume of the room to give priority to small rooms with a high occupancy, as quanta concentration builds up faster in small rooms.

As expected, applying good ventilation to less rooms instead of the entire school leads to less pronounced improvements. However, fig. 4-C shows that selecting only a fraction of rooms for improved ventilation, according to the criteria explained above, still results in median outbreak sizes comparable with those obtained in a setting with 30-40% vaccination coverage. In particular, the *size-corrected* strategy, which requires only information readily available to each school (i.e. school rosters and room size) can result in median outbreak sizes that are comparable to those obtained with vaccination rates above 40%, even when only applied to 25% of all rooms.

## Methods

### Data

The data used in this paper were collected at a US high school during one school day using wireless sensor technology. In total, 789 individuals (94% of the school population) participated. They wore small sensors that detect and record radio signals broadcast by other nearby sensors. Further, stationary devices broadcasting signals were attached to fixed locations (at least one per room) throughout the school campus to keep track of the participants’ locations.

Consequently, the data include two types of records. Close proximity interactions (CPIs) are records that indicate two participating individuals standing face-to-face with a distance of less than three meters at a certain point in time. Location records are records that indicate the presence of an individual nearby a stationary device (location information is at the level of rooms).

A detailed description on how information and noise were separated in the data is provided elsewhere^17^. Data were collected at time intervals of 20 seconds.

### Model of influenza spread

We used an individual-based model with a susceptible, exposed, infectious, recovered (SEIR)-type structure. We assumed that influenza is introduced into the school population by one index case at the beginning of a simulation run and that no further introductions from outside occur. The duration of a simulation time step was half a day. Individual *j*’ s probability *P*_*j*_ to switch from the susceptible to the exposed state depends on the mode of transmission. We defined one function *P*_*a,* *j*_ for aerosol transmission and one function *P*_*cc,* *j*_ for close-contact transmission, as laid out below. We ran simulations for an all-aerosol scenario, for an all-close-contact scenario, and one for a scenario where both aerosol and close-contact transmission occur as 0.5*P*_*a,* *j*_ + 0.5*P*_*cc,* *j*_. The duration of the exposed state follows a Weibull distribution with an offset of half a day; the power parameter is 2.21, the scale parameter is 1.10^11, 18^). After that period in the exposed state, every individual will be in the infectious state for exactly one time step before turning into home confinement and, finally, recovering. To allow for the fact that the onset of influenza symptoms is typically sudden and that affected individuals will be dismissed quickly, we reduced *P*_*j*_ by 75%, as described in^11^.

### Close-contact transmission probability

Assumptions, parameters, and structure of the close-contact transmission model are described in detail elsewhere^11^, and therefore only described briefly here.

Close-contact transmission require social interaction between an infectious and a susceptible individuals, and it includes transmission via large droplets that do not travel far and do not stay suspended in the indoor air as well as transmission via direct, physical contact^19^. Risk of transmission is usually operationalized as a function of contact duration^20^. Based on data from an outbreak on a commercial airliner^21^, the probability of transmission was estimated as

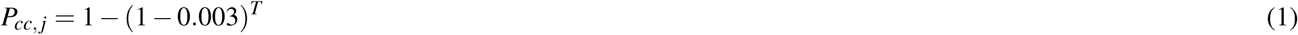

with *T* being the contact duration between two individuals in number of sensor recordings (every 20s)^11^.

### Aerosol transmission probability

In our model, we quantify amounts of aerosolized virus particles in ‘quanta’, following Wells’ ^22^ quantum theory of disease transmission. A quantum is defined as the amount of infectious droplet nuclei required to infect the fraction 1 *-* 1*/e* of a susceptible population exposed to it.

We assume that every room of the school is a well-mixed airspace that is only connected to the outside, but does not exchange air with other rooms. We further assume that removal of aerosolized virus particles by ventilation is the dominant removal process and that, e.g., neither inactivation nor settling play an important role. Under these assumptions, we model the concentration of virus particles in a particular room *r* as

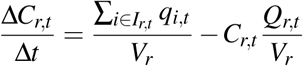

where *C*_*r,t*_ is the quanta concentration in room *r* at time *t*, *I*_*r,t*_ is the set of all infectious individuals that are in room *r* at time *t*, *q*_*i,t*_ is the quanta shedding rate of infector *i* at time *t*, *V*_*r*_ is the volume of room *r*, and *Q*_*r,t*_ is the fresh air supply rate of room *r* at time *t*. The quotient *Q*_*r,t*_ */V*_*r*_ is also known as the air change rate (ACR).

The instantaneous dose of infectious material, *D* _*j,t*_, inhaled by individual *j* at time step *t* is given by

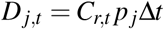

where *C*_*r,t*_ is the quanta concentration in room *r* - the room in which individual *j* is located at time *t* - at time *t*, *p* _*j*_ is the breathing rate of individual *j*, and Δ*t* is the duration of a simulation time step, here 20s.

Individual *j*’ s total exposure, *D* _*j*_ during an entire school day is given by

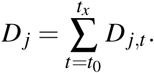

Combining the total daily exposure with Wells’ definition of quanta allows to model the probability *P*_*a,* *j*_ of a fully susceptible individual to become infected during one simulation school day as

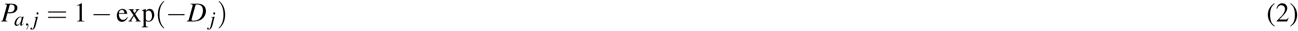

where the total exposure *D* _*j*_ is the only parameter required.

#### Shedding rate

Both bottom-up^23, 24^ and top-down approaches^25^ have been used by others to estimate quanta-based shedding rates for influenza. Bottom-up studies (mechanistic approaches starting from basic measurements and processes) suffered from huge uncertainties and differed by three orders of magnitude. We used data from Rudnick and Milton’ s^25^ top-down study that back-calculated quanta shedding rates from the same outbreak data^21^ that was also used to parameterize the close-contact model^11^. They estimated shedding rates of between 79 quanta/h and 128 quanta/h, depending on model assumptions. We chose a shedding rate of 100 quanta/h for our aerosol transmission model.

#### Ventilation rate

We assumed different scenarios for the ventilation rate. According to the ventilation recommendations for schools by the American Society of Heating, Refrigerating and Air Conditioning Engineers (ASHRAE)^13^, the ventilation rate for classrooms should be at least 8 l/s per person. Daisey et al.^26^ estimate that for a typical classroom situation, this corresponds to an air change rate (ACR) of 3.0 air changes per hour^26^. This estimate served as our good-ventilation scenario. Various studies found substantially lower ACR in US schools,^26–28^ and CO_2_ concentrations at the high school we collaborated with (unpublished data) indicated very poor ventilation conditions, too. In line with reported ACR in other US schools, we assumed 0.5 air changes per hour for a poor ventilation scenario.

#### Breathing rate

The breathing rate of humans depends mainly on their age, gender, and activity levels^29^. In line with Adams’ ^29^ measurements and in accordance with other work in the field^25^, we assumed a constant breathing rate of 8 l/min for every individual.

### Interventions

We compared ventilation-based interventions (effect only on aerosol transmission) with vaccinating individuals (effect independent on transmission pathway).

#### Ventilation rate

Baseline scenario to which all intervention scenarios were compared with was the poor ventilation scenario (ACR 0.5 h^−1^). Basic interventions were improving the ventilation to ASHRAE standards (ACR 3.0 h^−1^) and to achieve an intermediate improvement (ACR 1.5 h^−1^), respectively.

We further analyzed how ventilation improvement only in some rooms would affect infection spread. We defined three different methods to identify rooms for which the ACR was increased from 0.5 h^−1^ to 3.0 h^−1^: (i) *optimal*, using all available information from simulation runs, we identified rooms with the highest cumulative exposure, i.e., where most susceptibles will be exposed or where doses are highest in a typical simulation run; (ii) *schedule-based*, identifying rooms with the highest cumulative occupancy throughout a school day according to the school’ s official roster; (iii) *size-corrected*, which is similar to the schedule-based approach, but the total occupancy was divided by the volume of the room to give priority to small rooms with a high occupancy, as quanta concentration builds up faster in small rooms.

All three methods were used to identify rooms that represent 5%, 10%, 15%, 20% and 25% of the total indoor space. Methods (ii) and (iii) could realistically be applied in a school setting. Comparing interventions based on them with optimal ones allows assessing their relative performance to the theoretical optimum.

#### Vaccination

We assumed a vaccination effectiveness (protection against transmission) of 60% and a random distribution of vaccination status among the school population. We simulated the impact of vaccination for vaccination coverage values between 0% (baseline) and 100% with increments of 10%.

## Discussion

There is mounting evidence that aerosol-transmission is an important factor in the spread of influenza^3–9^. Despite this, virtually all infectious disease dynamics models on influenza have thus far ignored aerosol-transmission. Here, we parameterized a model with empirically obtained values to investigate the dynamics and control of influenza under the assumptions of no, partial, or full aerosol transmission. In order to create a realistic model, we used contact network and location data that was previously obtained at a US high school using wireless sensor network technology^11^. This dataset is well-suited for the objective of the study: modeling droplet-based transmission requires data on close proximity contacts, and modeling aerosol-transmission requires data on location in rooms and information about the rooms, such as the room size. The dataset used here, even though limited in scope, and particularly also in duration, contains all of this information.

Using empirical estimates of various influenza-related parameters, we found that the overall disease dynamics does not differ substantially between the models using no, partial, or full aerosol transmission. This isn’ t entirely surprising, given the fact that aerosol transmission parameters were estimated from the same influenza outbreak that was used to parameterize previous (pure droplet-based) influenza models. However, aerosol transmission does change the underlying transmission network (see fig. 1), which in turn could nevertheless have a substantial impact on disease dynamics, depending on the specific co-location patterns of individuals. Our finding that the dynamics do not change substantially in a model parameterized by empirical co-location data may simply be a reflection of the fact that schools are high density environments with comparatively limited movement, and that the underlying transmission-dependent network structures may be very similar.

While vaccination is at the heart of influenza prevention efforts, aerosol-based transmission of influenza opens up additional possibilities to control the spread of the disease. In particular, when infectious agents can remain airborne, air ventilation is a well known method to mitigate disease spread^14^. In this study, we assessed the effect of bringing the air change rates up to the recommended levels by the American Society of Heating, Refrigerating and Air Conditioning Engineers (ASHRAE)^13^, which defines an acceptable ventilation in classrooms to be 3 air changes per hour. We found that by doing so, we were able to generate reductions in expected outbreak sizes that would normally only be possible with a substantial vaccination coverage of 50-60%, which is within the range of observed vaccination rates in school settings^15^. Moreover, even when bringing only a quarter of the rooms to the recommended air change rates, using easy-to-obtain data in order to select the best rooms, we were still able to obtain outbreak size reductions that would require 30-40% vaccination coverage when air change rates are at levels commonly reported at US schools.

Our results suggest that improvements of ventilation in high density public spaces could be an important and easy-to-implement strategy supplementing vaccination efforts for effective control of influenza spread. This observation rests on the assumption that at a substantial part of influenza spread is due to aerosol-based transmission, for which there is mounting evidence. Given that increased air ventilation rates are not known to have any negative side effects, and that there are numerous infectious diseases that are entirely or partially transmitted via aerosol (e.g. tuberculosis), the findings here thus provide an additional argument corroborating the public health recommendations for good air ventilation. It should be noted that influenza vaccine effectiveness is often less than the here assumed 60%^16^, whereas good ventilation would reliably provide increased protection, further underlining its importance.

## Acknowledgements

This research was supported by a fellowship from the German Academic Exchange Service DAAD to T.S. (Grant D/10/52328); T.S. also thanks the UK National Institute for Health Research Health Protection Research Unit (NIHR HPRU) in Modelling Methodology at Imperial College London in partnership with Public Health England (PHE) for funding (grant HPRU-2012- 10080). The views expressed are those of the authors and not necessarily those of the MRC, the NHS, the NIHR, the Department of Health, Public Health England or any other organization the authors’ are affiliated with or received funding from. We also thank the High-Performance Computing group (SCITAS) of EPFL for his technical support.

## Author contributions statement

T.S. and M.S. conceived and designed the study. M.S. collected the data. G.L. and T.S. performed the simulations and analyses. All authors wrote, read and approved the final manuscript.

